# Seumetry: a versatile and comprehensive R toolkit to accelerate high-dimensional flow and mass cytometry data analysis

**DOI:** 10.1101/2024.07.23.604747

**Authors:** Malte Borggrewe, Markus Flosbach, Hamburg Intestinal Tissue Study Group, Stefan Bonn, Madeleine J. Bunders

## Abstract

Recent progress in flow and mass cytometry technologies enables the simultaneous measurement of over 50 parameters for an individual cell. The resulting increase in data volume and complexity present challenges, as conventional analysis methods based on manual gating are time-consuming and fail to capture unknown or minor cell populations. Advances in single-cell RNA sequencing (scRNAseq) technologies have prompted the development of sophisticated computational analysis tools specifically designed to process and analyze high-dimensional biological data, some of which could significantly improve certain aspects of cytometry data analysis. Building on these advances, we here present Seumetry, a framework that combines flow and mass cytometry data-specific analysis methods with the capabilities of Seurat, a powerful tool for the analysis of scRNAseq data. Seumetry offers advanced quality control, data visualizations, and differential population abundance and protein expression analysis. We tested Seumetry on an in-house generated complex dataset of immune cells from different layers of human intestines, demonstrating that Seumetry accurately identifies distinct immune cell populations. Furthermore, using a publicly available mass cytometry dataset, Seumetry recapitulates previously published results, further validating its use for high-dimensional flow and mass cytometry data. In summary, Seumetry provides a new scalable framework for the comprehensive analysis of high-dimensional cytometry data with seamless integration into commonly used scRNAseq analysis tools, enabling in-depth analysis methods to facilitate biological interpretations.

## Background

Advances in flow and mass cytometry technologies now enable the simultaneous measurement of over 50 parameters per cell in millions of cells per sample. This high-dimensional data poses new demands to existing analysis techniques. Conventional cytometry analysis based on 2-dimensional manual gating, until now the gold standard for cytometry data analysis, faces challenges in scaling for the application of high-dimensional cytometry data. Issues with conventional gating strategies include inefficiency, subjectivity, and a significant risk of missing unknown or small cell populations. Hence, improved and unbiased computational methods are needed to harness the potential of high-dimensional cytometry data.

To date, several computational tools for cytometry data analysis have been developed, primarily designed for the R programming language, including CyTOF workflow^1^, diffcyt^2^, FlowSOM^3^, flowCore^4^, flowAI^5^, flowViz^6^, CytoTree^7^, CITRUS^8^, ImmunoCluster^9^, flowWorkspace^10^, and PeacoQC^11^. These individual workflows address specific aspects of high-dimensional cytometry data analysis, including handling flow cytometry standard (FCS) files, preprocessing, dimensionality reduction, clustering based on self-organizing maps, and differential abundance analysis. However, interoperability between these packages is limited and inefficient, and methods for advanced data analysis such as data integration and trajectory analysis are scarce^12–14^.

In contrast to the lack of high-dimensional cytometry data analysis methods, algorithms for analyzing single-cell RNA sequencing (scRNAseq) data are highly developed and offer a wide range of analysis possibilities. The most commonly used R framework for scRNAseq data analysis is Seurat^15^, which covers every aspect of data analysis from preprocessing to a range of downstream analyses, including dimensionality reduction and clustering. A notable advantage of scRNAseq tools is their interoperability, since most third-party tools that offer additional analysis options such as batch correction^15,16^ or trajectory analysis^17,18^, integrate seamlessly into the Seurat workflow. However, these scRNAseq analysis algorithms are not readily accessible for the analysis of cytometry data. With these possibilities and challenges in mind, we developed the R toolkit Seumetry, a flow and mass cytometry analysis framework that offers state-of-the-art analysis options for comprehensive cytometry data analysis. Seumetry includes a broad range of cytometry-specific algorithms for data handling, while it seamlessly integrates with Seurat, providing access to the latest scRNAseq analysis options.

## Overview of the Seumetry workflow

Currently available flow and mass cytometry analysis tools focus on specific aspects of data analysis, whereas Seumetry covers all aspects of cytometry data (pre)processing and analysis (Table 1). Furthermore, Seumetry offers integration into the Seurat framework to access advanced scRNAseq algorithms and unique features such as the detection of antibody aggregates, which are associated with complex antibody panels required for high dimensional cytometry (Table 1).

**Table 1.**
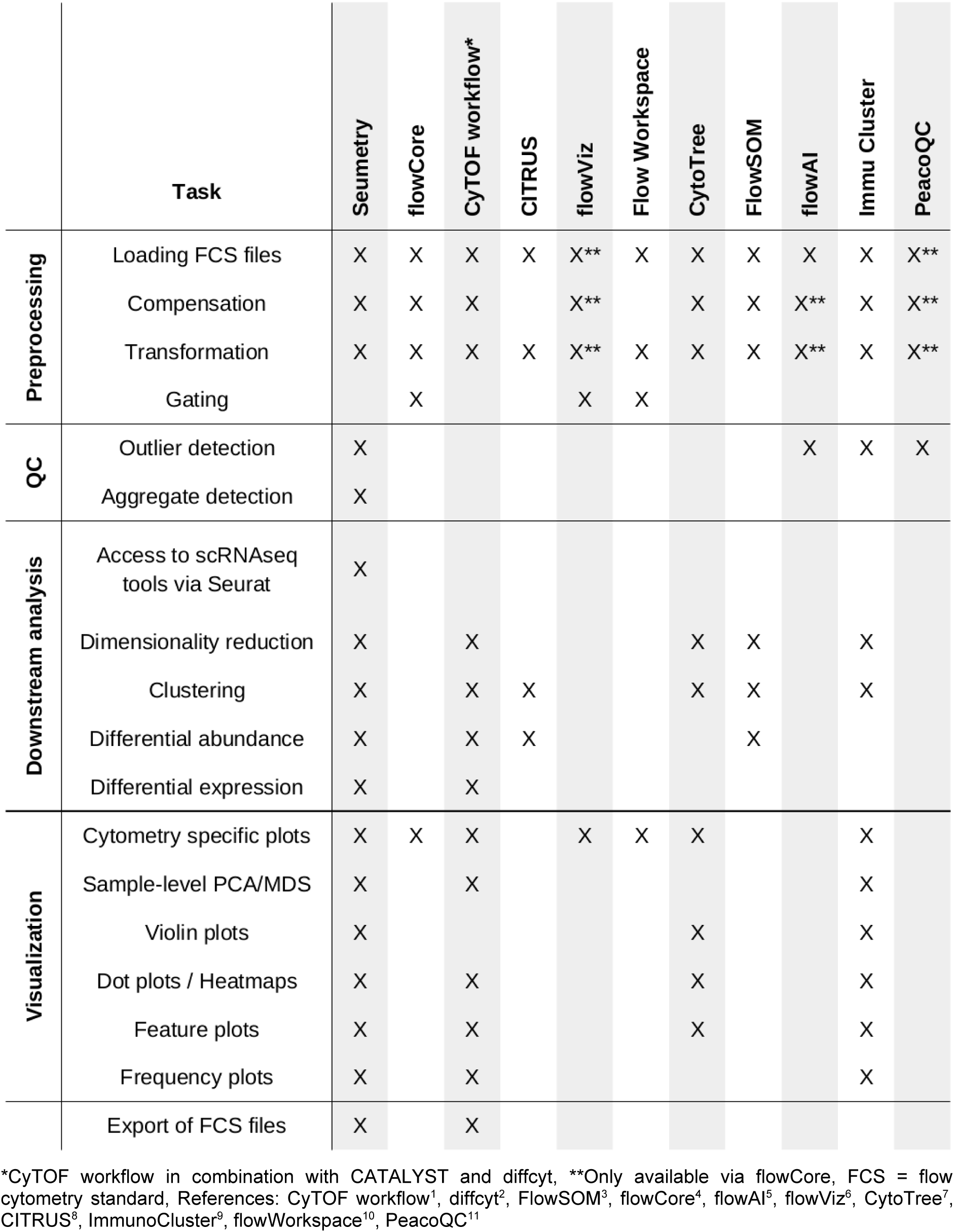
Comparison of Seumetry to other R packages for cytometry data analysis.

To demonstrate the functionality and workflow of Seumetry, we generated a high-dimensional spectral flow cytometry dataset comprising 39 protein markers of human intestinal immune cells (Table S1 and S2). The test dataset consists of immune cells isolated from 7 adult human intestines, separated in epithelium and lamina propria layers (Fig. 1). The data was manually pre-gated on single (FSC, SSC) live (viability dye) immune (CD45 positive) cells and downsampled to a maximum of 20,000 cells per sample. This dataset is used to present all functionalities of Seumetry with the aim of annotating intestinal immune cell populations and assessing differences in epithelial and lamina propria-derived immune cells.

**Figure 1.**
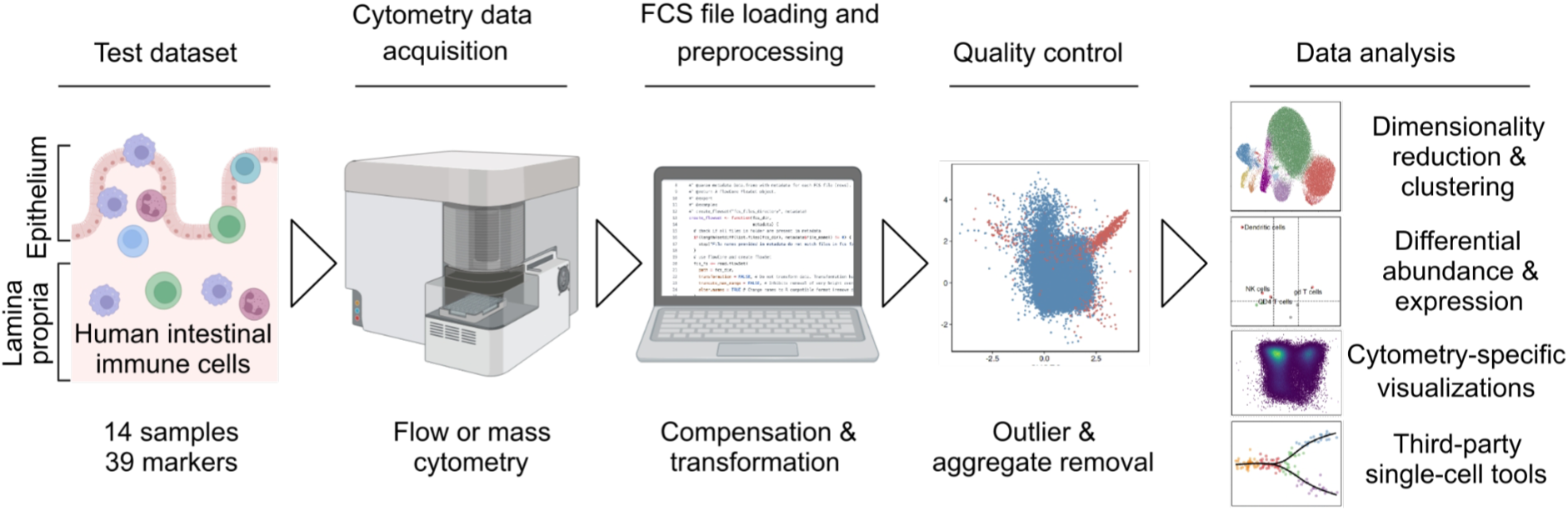
Generation of flow cytometry test dataset and Seumetry workflow. (A) To showcase the Seumetry workflow, we generated spectral flow cytometry data of immune cells in intestinal mucosal samples from 7 adult donors, including separate analysis of epithelial layer and lamina propria-derived cells. Cytometry raw data is loaded into R and preprocessed, including compensation of protein channels and data transformation. Quality control methods include the detection and removal of outlier events and antibody aggregates. Data analysis includes dimensionality reduction and clustering, differential abundance, and differential expression analysis, additional visualization methods, and potential application of third-party single-cell tools.

After flow cytometry standard (FCS) file loading, data is preprocessed in Seumetry including compensation and transformation (Fig. 1). Since cytometry data can be inherently noisy, we implemented multiple algorithms to improve data quality, including the identification and removal of outliers and antibody aggregate artifacts. Antibody aggregates can occur in highly complex cytometry panels which include a large number of antibodies that can form aggregates and lead to false positive signals^19^. Seumetry contains a novel algorithm for the automated and unbiased removal of these antibody aggregates.

Finally, the high-quality data can be analyzed using common scRNAseq algorithms via integration into the Seurat workflow including dimensionality reduction, clustering, batch correction, and trajectory analyses^15–18^. To expand analysis options, we also implemented additional visualizations and methods for differential abundance and expression analysis (Fig. 1).

Using Seumetry to analyze the in-house generated flow cytometry-based dataset of intestinal immune cells, and a publicly available dataset of immune cells analyzed by mass cytometry, we conclude that Seumetry can be used for comprehensive analysis of flow and mass cytometry data and accelerate biological interpretation of complex cytometry data.

## Data preprocessing and removal of outliers

Seumetry incorporates all necessary flow and mass cytometry-specific data preprocessing and quality control steps. For cytometry data, compensation may be necessary to correct for spillover effects of protein channels, which is included in Seumetry using the compensation matrices embedded in the FCS files or external matrices (Fig S1A). Next, Seumetry offers cytometry-specific options for data normalization and transformation to stabilize variance and convert data from an exponential to a linear scale. Transformations include biexponential for flow cytometry and the more commonly used inverse hyperbolic sine function (arcsinh) for flow and mass cytometry (Fig. S1B).

To eliminate undesired outliers in the data, which commonly manifest as very low or high-intensity events in flow and mass cytometry data, e.g. resulting from cell debris, dead cells, or cell clumps, Seumetry offers both manual and automatic workflows. In the manual workflow, users can visually inspect each parameter and provide positive and negative thresholds for each channel. Subsequently, events above or below the specified thresholds are removed from the data. The automatic workflow employs an isolation forest, which calculates an outlier score for each event based on all provided antigen channels. Events are regarded as outliers if they surpass a specified outlier score threshold (default: 0.7) and can subsequently be removed to obtain high-quality data. To test the performance of the automatic outlier removal workflow, we assessed ungated data from our generated intestinal immune cells dataset containing noisy events, such as apoptotic cells and cell clumps (n=3; downsampled to 500,000 cells per sample). Although the isolation forest only used measurements of antigen channels, our automatic workflow almost exclusively detected events with unexpectedly high measurements for cell size and granularity (cell clumps), or dead cells (Fig. 2A), indicating that the algorithm removes noisy events while keeping high-quality cells. Subsequently, the algorithm was used to improve the quality of the pre-gated intestinal immune cell test dataset, which resulted in the removal of 60 events with very high or low protein signal intensity (Fig. 2B). Together, this automatic outlier removal method is computationally efficient and can detect noisy events while retaining high-quality cells.

**Figure 2.**
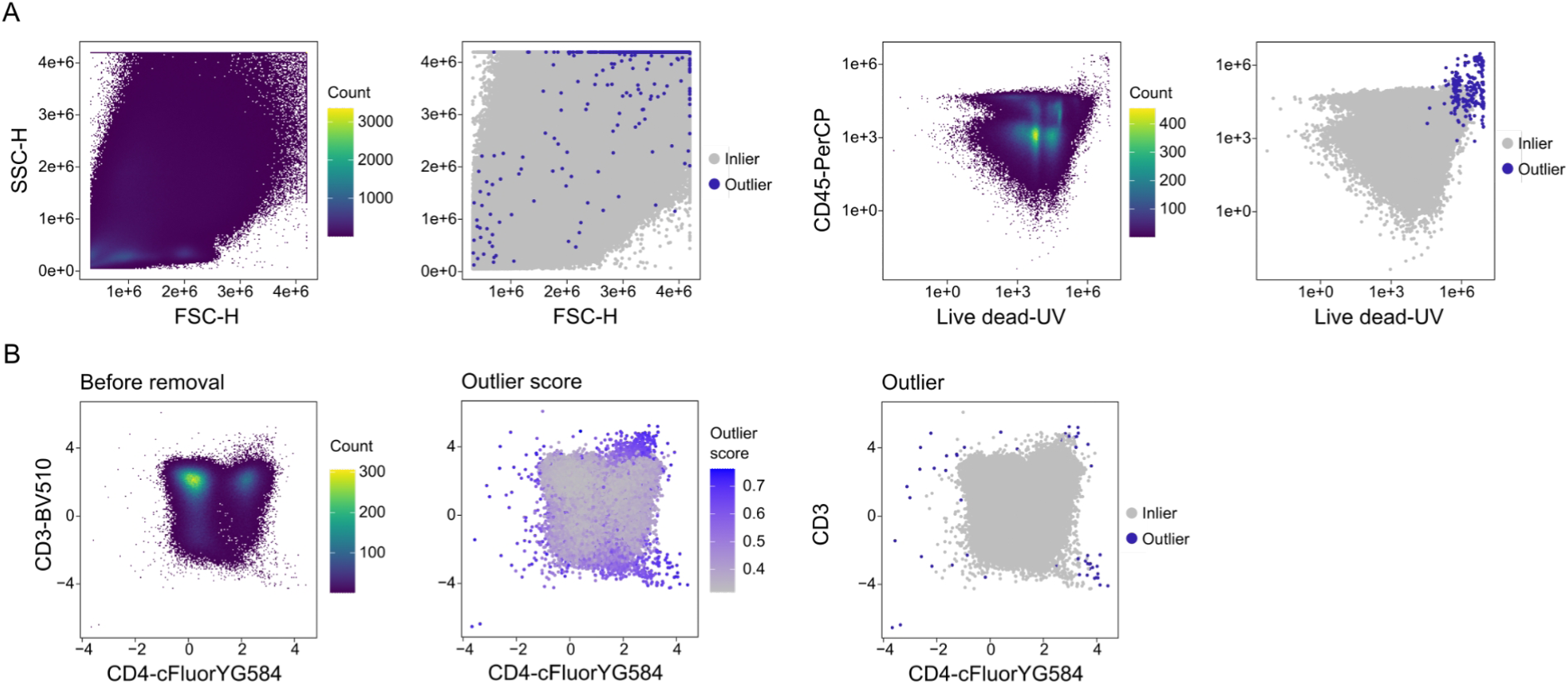
Detection and removal of outlier events in flow cytometry data. (A) Two-dimensional density plots and scatter plots illustrating outliers (blue) and inliers (gray) of ungated events showing SSC-H vs FSC-H and CD45-PerCP vs live dead (n=3). (B) Representative plots of outlier detection and removal in test data using CD3 and CD4 as examples (n=14).

## Detection and removal of antibody aggregates

The use of large antibody panels in high-dimensional cytometry can introduce additional artifacts due to the aggregation of two or more antibodies, leading to false positive signals on cells (Fig. 1: Quality control). Adjusting laboratory protocols to individually stain cells for each antibody can mitigate these artifacts. However, this approach can be highly time-consuming for large antibody panels, posing new challenges affecting cell viability and staining stability. Alternatively, antibody aggregates can be removed using manual gating, which is time-consuming and subjective, introducing observer variability. To provide an unbiased approach for removing antibody aggregates, we developed a novel algorithm to detect aggregate signals on cells.

In the initial step, the algorithm identifies antigen channels that potentially contain signals of antibody aggregates. Signals of antibody aggregates are expected to be highly collinear double-positive events in specific antigen channel combinations. Thus, the data is subset to double-positive events (default threshold > 1), and a Pearson correlation is performed (Fig. 3A). A high Pearson R value may suggest the presence of highly collinear events, indicative of cells with antibody aggregates. Applying a threshold for Pearson R (default: 0.7) identifies antigen channel combinations with potential aggregates (Fig. 3B), which resulted in 16 channel combinations with a high likelihood of antibody aggregates in our dataset (Fig. 3C and Fig. S2).

**Figure 3.**
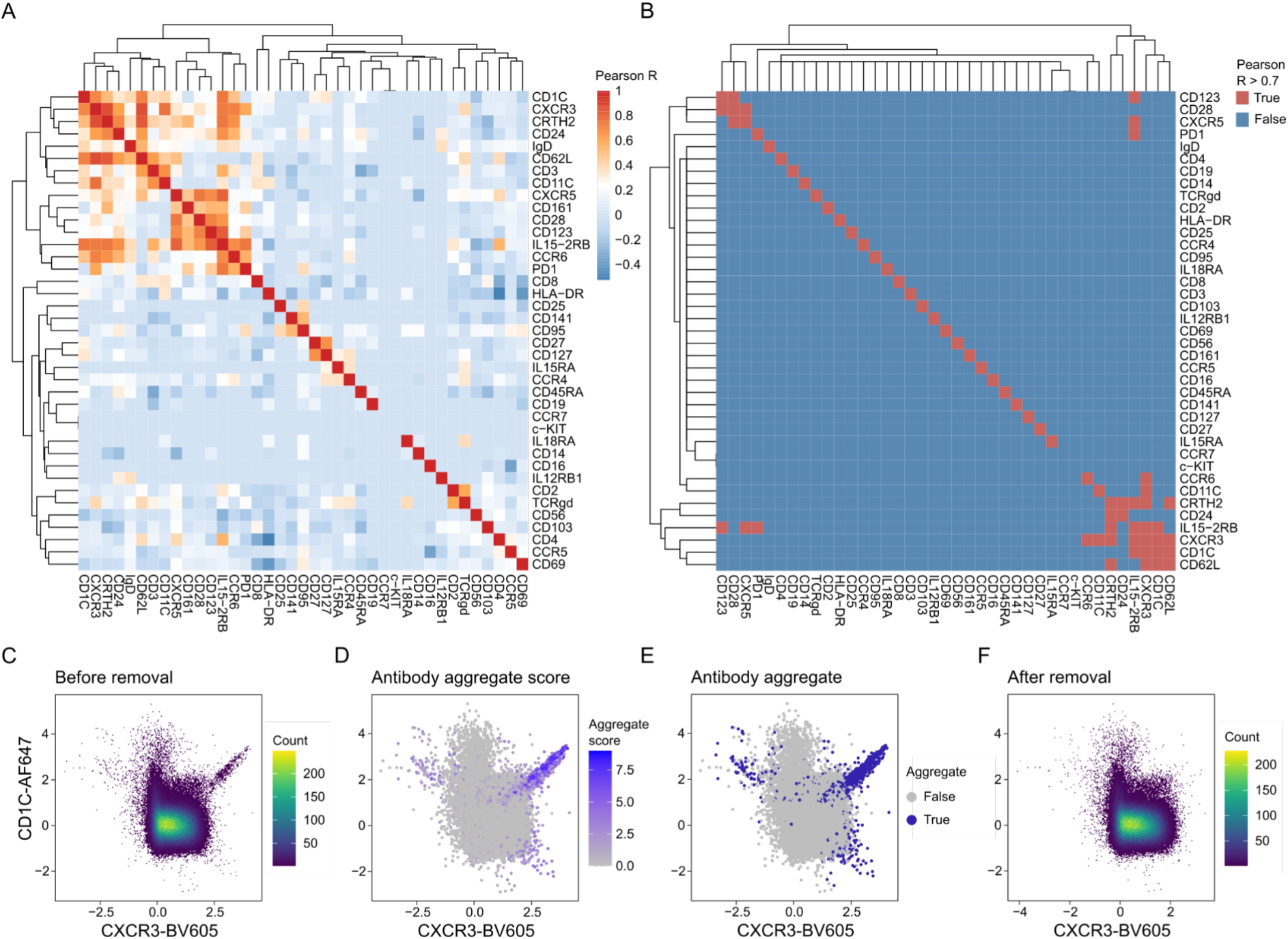
Detection and removal of antibody aggregates. (A and B) Correlation heatmap depicting (A) Pearson R and (B) Pearson R > 0.7 for double-positive events of each channel combination. (C-F) Representative plots of antibody aggregate detection and removal using CD1C and CXCR3 as examples. (C) Two-dimensional density plot of events before removal of antibody aggregate. (D) Scatterplot visualizing aggregate score for each event. (E) Scatterplot depicting antibody aggregates (blue) and non-aggregates (gray) based on an antibody aggregate score threshold of 2. (F) Two-dimensional density plot of events after removal of antibody aggregates (n=14).

To identify potential antibody aggregates in the above detected channel combinations, we developed a modified random sample consensus (RANSAC) algorithm. RANSAC is an iterative method for fitting a linear model in the presence of outliers^20^, which prevents the modeling of non-aggregate double-positive cell populations. We modified the algorithm to utilize the upper 50% of positive events with a slope between 0.75 and 1.25 to model only highly collinear antibody aggregate events. Events with a low residual value (default residual value: less than 0.25) based on the fitted linear model are labeled as potential antibody aggregates. The antibody aggregate score indicates the number of channel combinations in which an event is labeled as a potential aggregate (Fig. 3D). To prevent the removal of non-aggregates, events are only considered antibody aggregates if they are labeled as potential aggregates in two or more channels by default (Fig. 3E and Fig. S2), and these events are subsequently removed (Fig. 3F). Detection of outliers based on isolation forest (see above) can additionally be performed after removal of aggregates to filter out remaining outliers.

In summary, our algorithm for automatic antibody aggregate detection offers an efficient and unbiased approach to removing antibody aggregates, generating improved quality in cytometry datasets.

## Dimensionality reduction and clustering of intestinal immune cell populations

Many computational algorithms have been developed to analyze scRNAseq data, which are not easily accessible for high-dimensional cytometry analysis^14^. These tools include common algorithms for dimensionality reduction and clustering and more specialized algorithms such as trajectory analyses and batch correction^15–18^. We developed Seumetry as a pipeline for researchers to now seamlessly employ a multitude of scRNAseq algorithms for the in-depth analysis of the flow and mass cytometry data.

To explore the variance between samples, for example, to assess the quality of replicates and differences between conditions, we added a principal component analysis (PCA) based on median protein expression per group (Fig. 4A). As our dataset consisted of human samples that exhibited considerable donor variation (Fig. 4A), the data was integrated using an anchor-based CCA integration supplied in the Seurat framework^15^. Subsequently, a uniform manifold approximation and projection (UMAP) and Louvain clusters were computed based on corrected PCs (Fig. S3A). To identify specific intestinal immune cell populations, differential cluster markers were calculated (Seurat “FindAllMarkers”) (Fig. S3B), and clusters were merged and annotated (Fig. 4B). The distribution of immune cell populations was stable across donors (Fig. 4C), demonstrating a successful donor integration. This analysis workflow resulted in the identification of the expected major intestinal immune cell populations with the detection of their known markers (Fig. 4D).

**Figure 4.**
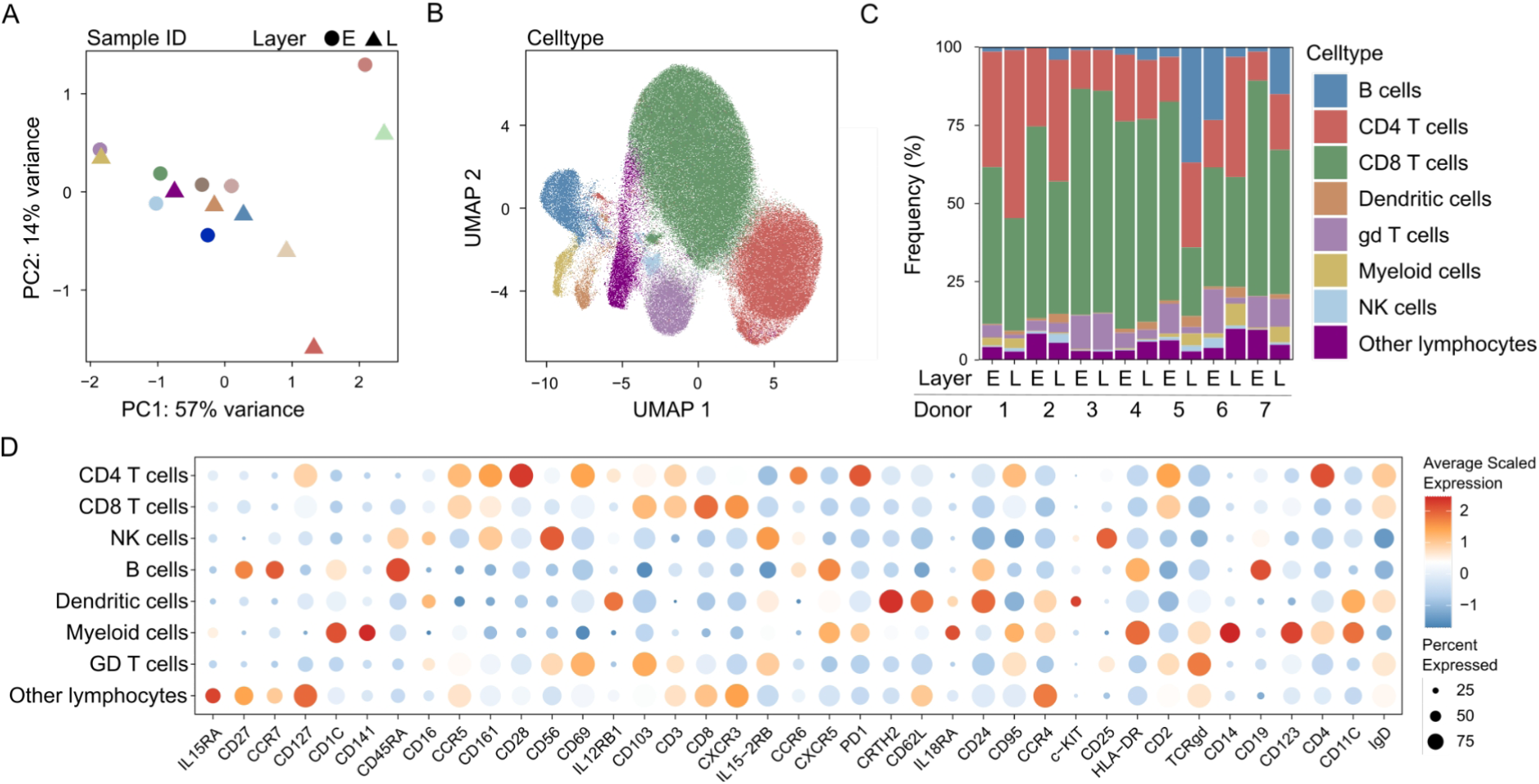
Dimensionality reduction and clustering of intestinal immune cells. (A) Principal component analysis based on median surface protein expression per sample, colored by sample id and shape indicates layer. (B) UMAP of all samples colored by immune cell populations identified by merging and annotating louvain clusters. (C) Frequency plot of immune cell populations across all samples. (D) Average expression (scaled) of all proteins across immune cell populations visualized in a dotplot (n=14). E = Epithelium, L = Lamina propria

## Differential population abundance and protein expression of intestinal immune cells in epithelium and lamina propria

High-dimensional cytometry is commonly used to identify the differential abundance of cell populations or differential expression of proteins across conditions. Hence, we implemented a differential abundance and expression analysis.

Differential abundance analysis in Seumetry utilizes cell numbers per sample and condition to construct a generalized linear model (GLM) using the edgeR R package^21^. Pairwise comparisons can be performed, which uses a likelihood ratio test and provides corrected p values using the Benjamini-Hochberg method. To demonstrate the differential abundance analysis, we compared the abundance of immune cells between the epithelium and lamina propria layers (Fig. 5A and B). Dendritic cells, NK cells, and CD4 T cells were enriched in the lamina propria, whereas γδ T cells were enriched in the epithelial layer of the intestines. These findings confirm previous findings by our group and others^22–24^. In line with these studies, CD8 T cells were higher in the epithelium (Fig. 5B), albeit not significant (adjusted p = 0.15), further validating that Seumetry accurately identifies immune cell populations in an unbiased manner.

**Figure 5.**
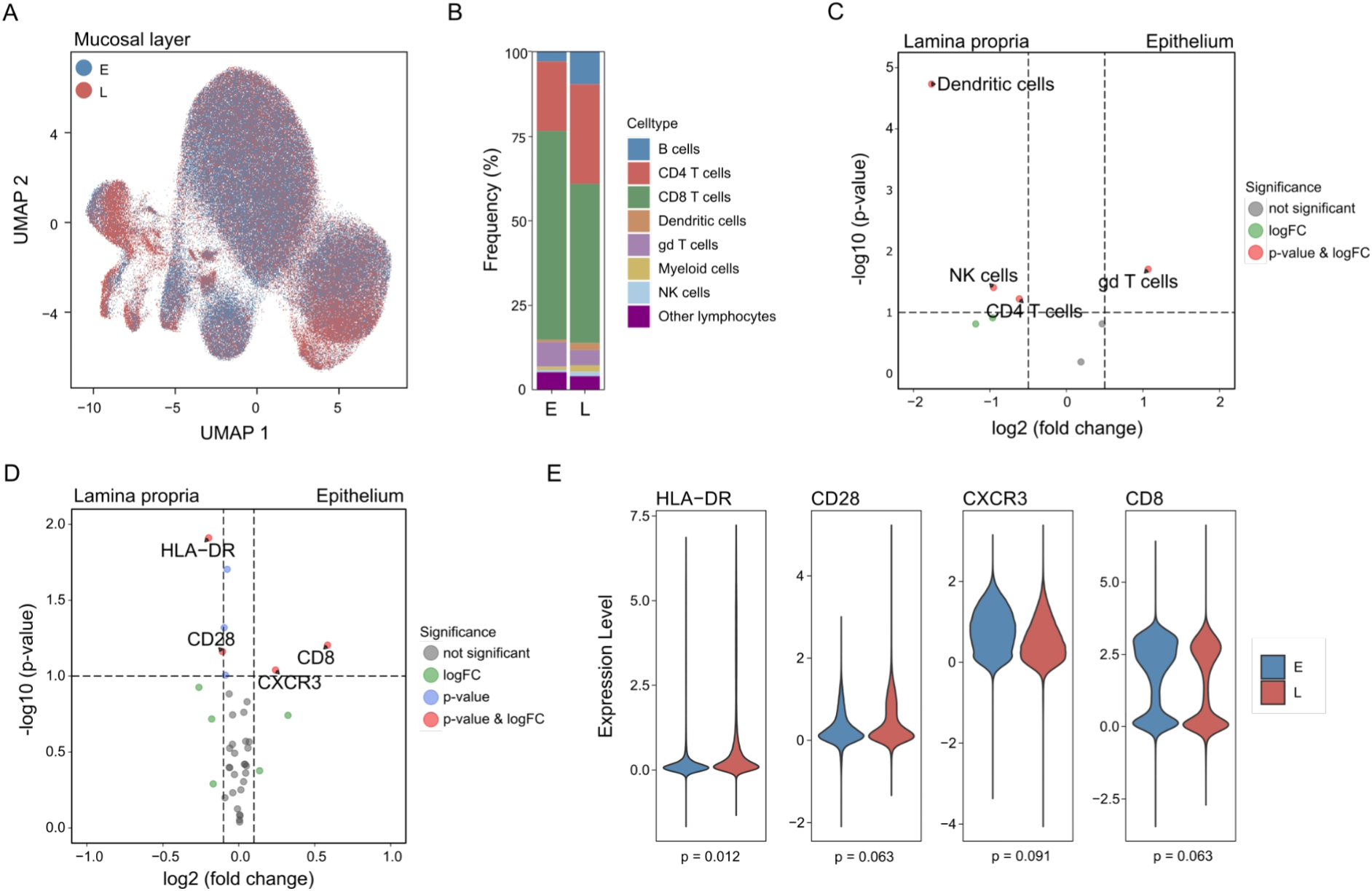
Differential abundance and differential expression analysis of immune cells in intestinal mucosal layers. (A) UMAP of all samples colored by mucosal anatomical layers. (B) Frequency plot of immune cell populations in epithelium (E) and lamina propria (L). (C) Volcano plot of differential abundance analysis comparing immune cell populations in epithelium to lamina propria (FDR < 0.1, logFC > 0.5). (D) Volcano plot of differential expression analysis using median protein signal intensity per sample comparing immune cells of epithelium to lamina propria (FDR < 0.1, logFC > 0.1). (E) Violin plot of differential protein expression across mucosal layers (n=14).

To investigate differences in (surface) protein expression in cytometry data, a differential expression analysis can be conducted on a single-cell level using Seurat’s ‘FindMarkers’ function; however, using the median protein expression per sample (pseudobulk) provides a more conventional approach to detect differentially expressed markers in cytometry. To this end, we used the limma R package^25^ to compare the median protein signal intensity per sample across conditions. The function computes a linear model followed by an empirical Bayes statistic for differential expression (eBayes). HLA-DR and CD28 were enriched on immune cells in lamina propria samples, whereas CXCR3 and CD8 were enriched on epithelial immune cells (Fig. 5D and E). HLA-DR was present on B cells (Fig. 4D), which are known to be enriched in lymphoid follicles in the lamina propria compared to epithelial layer in intestines^22^, whereas CXCR3 was high on CD8 T cells (Fig. 4D) and enriched in epithelial immune cells (Fig. 5D and E). In sum, the differential protein expression results are in line with the differential population abundance analysis and previous studies^22–24^. These findings highlight that the Seumetry toolkit offers a user-friendly implementation of a more conventional differential population abundance and differential protein expression analysis.

## Verification of Seumetry workflow using a publicly available dataset

To further demonstrate the fidelity and straightforward usability of Seumetry, we analyzed a publicly available dataset previously used to evaluate the cytometry analysis packages CyTOF workflow^1^ and CITRUS^8^. This mass cytometry dataset contains peripheral blood mononuclear cells (PBMCs) from two treatment conditions (control and BCR/FcR-XL-treated) stained for 10 surface and 14 intracellular antigens^1,8,26^. To be able to compare results, we used a similar approach as in the CyTOF workflow^1^, including downsampling of data to 1,000 cells per sample and using the 10 surface markers directly to compute the UMAP^1^. Furthermore, an over-clustering approach was used and the clusters were subsequently manually merged based on cell type^1^.

To explore the variance across samples in the public dataset, a PCA computed with Seumetry revealed a segregation of samples based on treatment conditions (Fig 6A), consistent with published results^1^. Seumetry clustering analysis yielded 20 clusters (Fig. 6B), which were manually merged based on the expression of the immune cell lineage markers (Fig. 6C and Fig. S4A). The resulting merged clusters exhibited similar lineage marker expression and frequencies of immune cell populations compared to published results from CyTOF workflow^1^ (Fig. 6D), with the identification of an additional CD45+ subset (Fig. 6C and D). Frequencies of immune cell populations per condition (Fig. 6E) and per sample (Fig. S4B) identified by Seumetry were similar to the published results^1^ and the differential abundance analysis revealed comparable results (Fig. 6F). In both, Seumetry and CyTOF workflow results^1^, NK cells and CD8 T cells were increased in treated compared to control samples, whereas B cells and CD4 T cells were decreased (Fig. 6F). Differentially expressed protein analysis based on Seumetry also aligned well with the published results^1^, with markers such as pBtk, pAkt, pErk, and pS6 being significantly differentially expressed in treated compared to control in both analyses (Fig. 6G).

**Figure 6.**
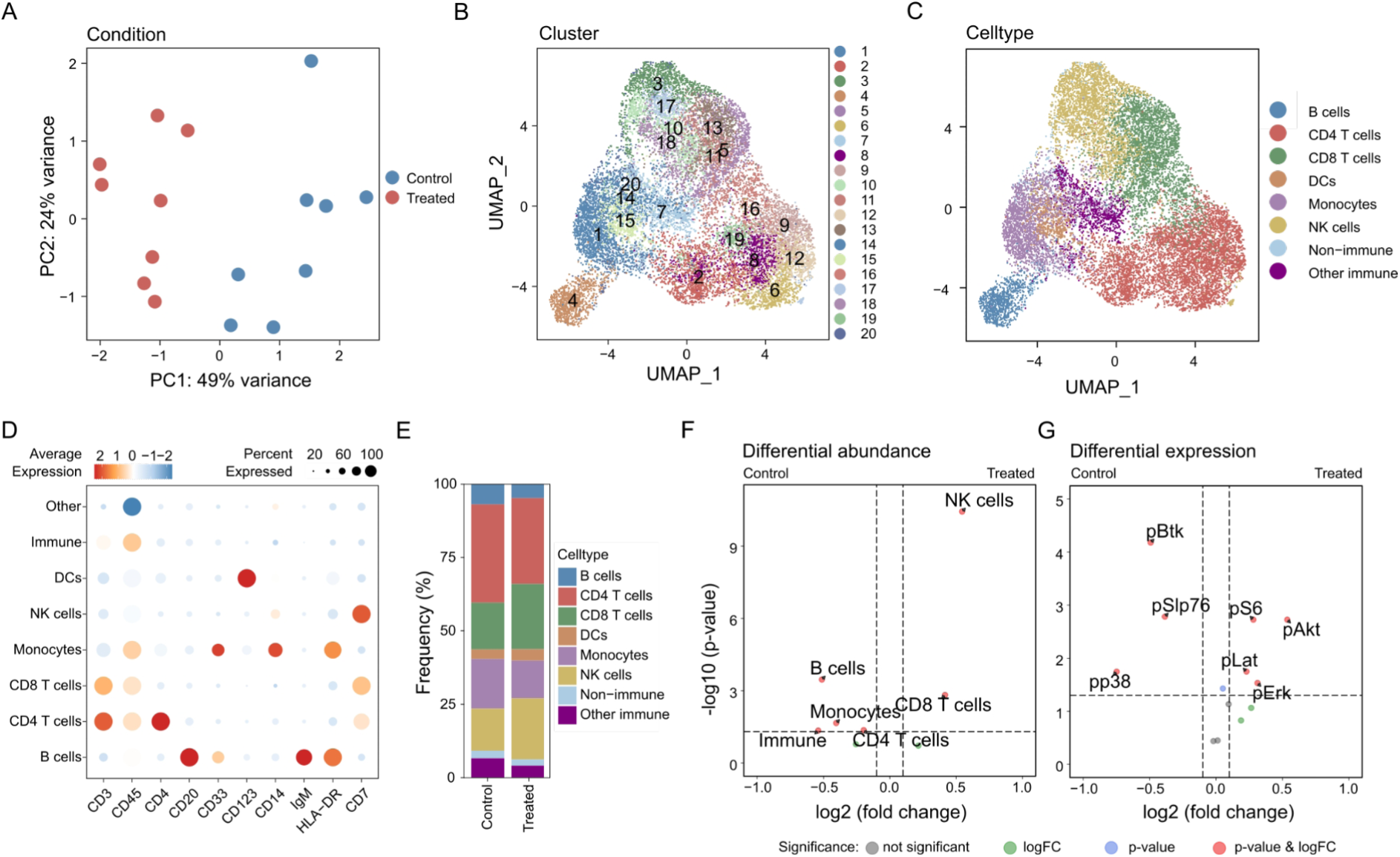
Analysis of publicly available mass cytometry data using Seumetry. (A) Principal component analysis based on median protein expression per sample colored by condition. (B-C) UMAP of all samples colored by (B) the 20 Louvain clusters and (C) merged clusters based on protein expression. (D) Dotplot of average lineage marker expression (scaled) in merged clusters. (E) Frequency plot of clusters across conditions. (G) Volcano plot of differential abundance analysis comparing stimulated to control (FDR < 0.05, logFC > 0.1). (H) Volcano plot of differential expression analysis using median protein signal intensity per sample comparing treated to control (FDR < 0.05, logFC > 0.1).

In conclusion, Seumetry recapitulates known and published differences between treated and control PBMCs^1,2^, underscoring its applicability, robustness, and reliability.

## Concluding remarks

Here, we introduce Seumetry, a versatile R toolkit for comprehensive analysis of high-dimensional cytometry data. Seumetry comprises all necessary tools to analyze flow and mass cytometry data: 1) preprocessing (compensation, transformation), 2) quality control (outlier and antibody aggregate detection), 3) cytometry-specific visualizations, 4) differential abundance and expression analysis. Furthermore, Seumetry offers advanced high-dimensional analysis methods such as dimensionality reduction, clustering, batch correction, and compatibility with third-party scRNAseq tools through seamless integration into the Seurat framework.

The rapid development of cytometry technologies allows us to quantify more protein markers. With the increasing complexity of cytometry data, traditional 2-dimensional manual gating methods are becoming less suited, as they are highly time-consuming, and due to the increased likelihood of missing cell populations. These issues with manual gating can only be alleviated using more automated approaches, which Seumetry provides. Automated analysis methods could facilitate high-dimensional cytometry data analysis and accelerate basic and translational research. As immunological analyses for clinical purposes are steadily increasing due to the enhanced use of biologicals and small molecules targeting immune pathways, standardized approaches such as Seumetry can aid robust and efficient immunological analyses for diagnosis and treatment monitoring.

In the future, the number of parameters will likely increase which will further exacerbate the challenges facing manual gating. In contrast, increasing data complexity can improve the unbiased analysis of cytometry data using Seumetry, since more information will be available to classify cell types and cell phenotypes. Seumetry is highly scalable, due to its interoperability with efficient scRNAseq toolkits, and methods and toolkits to analyze scRNAseq data are continuously evolving and improving, which will also be beneficial for cytometry data analysis using Seumetry.

One of the biggest challenges with cytometry data is the presence of noise, due to cell debris/clumps, apoptotic cells, or other artifacts such as antibody aggregates^10,11,19^. Preprocessing in Seumetry improves data quality by detecting noisy events in an unbiased manner. However, data quality is highly dependent on the source of cells/tissue and the processing in the laboratory. To alleviate these issues, we included multiple parameters that researchers can use to tailor the algorithm to each dataset, which should facilitate data-specific detection of noisy events based on scientific requirements.

High-dimensional cytometry and scRNAseq generate similar types of data on single-cell level, but are based on distinct modalities (protein and mRNA) that are highly complementary. Compared to scRNAseq technologies, which offer genome-wide information, cytometry is only able to capture a limited number of markers. However, cytometry measures proteins, which are a better predictor of biological function than mRNA expression, and cytometry is easily accessible and cheaper than scRNAseq. Future applications include the integration of scRNAseq and cytometry results to facilitate multi-omics analyses and address biological questions at different levels of cellular regulation. Furthermore, methods for imputation of protein expression in cytometry data with limited protein marker panels based on scRNAseq datasets will be interesting to explore. Seumetry can facilitate the development of novel analysis methods based on scRNAseq datasets through its compatibility with most scRNAseq methods and frameworks.

In summary, Seumetry offers a versatile and scalable R framework for the unbiased and comprehensive computational analysis of high-dimensional flow and mass cytometry data, which will be valuable for extracting meaningful biological information and facilitating the development of novel methods for cytometry data analysis.

## Materials & Methods

### Acquisition and processing of human intestinal tissue

Intestinal human tissues were collected upon surgical interventions for the treatment of intestinal diseases at the University Medical Center Hamburg-Eppendorf. Donors provided written informed consent and the collection and use of tissue samples were approved by the ethics committee of the Medical Association of the Freie Hansestadt Hamburg (Ärztekammer Hamburg). For an overview of donors please refer to supplementary table S1. Human intestinal tissues were processed and immune cells were isolated as described previously^23,27^. After mechanical removal of fat and muscular layers, tissues were incubated with Iscove’s modified Dulbecco’s medium (Thermo Fisher Scientific, 10135083) supplemented with EDTA (5 mM, Promega, V4231), 1,4-dithiothreitol (Carl-Roth GmbH+Co. KG, 6908.1), and 1% fetal bovine serum (Capricorn, FBS-11A) to obtain epithelial immune cells. The cell suspension was filtered through a 70-um cell strainer and intraepithelial lymphocytes were enriched after epithelial cell layer dissociation using density gradient centrifugation (LSM-A, Capricorn). Lamina propria immune cells were isolated by mincing and incubating in Iscove’s modified Dulbecco’s medium supplemented with 1 mg/mL Collagenase D (Sigma-Aldrich, 11088882001), 1% FBS, and 1000 U/mL DNaseI (StemCell Technologies, 07470). The resulting cell suspension was filtered using a 70 μm cell strainer and immune cells were obtained by density gradient centrifugation using 60% standard isotonic Percoll (Sigma-Aldrich, 17-0891-62).

### Flow cytometry

Cells were stained for 40 markers using a multi-step staining protocol. For a list of all antibodies and steps, see supplementary table S3. Cells were thawed and incubated in Hank’s balanced salt solution (Sigma, H-6648) supplemented with 0.1mg/ml DNAse (Sigma, D-4513) and magnesium chloride hexahydrate (Sigma, M-2670). Subsequently, cells were incubated in staining step 1 (Table S3) with the live dead UV blue dye 10 min at 4°C, followed by incubation in Human TruStain FcX Fc Receptor Blocking Solution (Biolegend, 422302) for 10 min at 4°C in FACS Buffer (PBS supplemented with 1% FBS). After washing with PBS, cells were incubated with antibodies of staining step 2 (Table S3) in FACS Buffer for 30 min at 37°C. Finally, antibodies of staining step 3 (Table S3) were added to the cells in FACSBuffer for 30 min at room temperature. Cells were washed in FACSBuffer and analyzed on Cytek Aurora. Before processing with the analysis, raw data was unmixed and cells were pre-gated on single cells (FSC, SSC) live (live dead UV negative) immune (CD45 positive) cells.

### Availability of code and data

Seumetry code is open source under an MIT license and can be viewed and used via Github: https://github.com/imsb-uke/Seumetry

Flow cytometry data generated for this manuscript is available via Zenodo: https://doi.org/10.5281/zenodo.11935872

## Acknowledgments

We thank all intestinal tissue donors who participated in this study and the Department of General, Visceral, and Thoracic Surgery of the University Hospital Hamburg-Eppendorf for the collection of intestinal tissues. The Hamburg Intestinal Tissue Study Group members include: Felix J. Flomm, Sebastian Schloer (Department of Virus Immunology, Leibniz Institute of Virology, Hamburg, Germany); Alaa Akar, Cornelius Flemming, Niklas Jeromin, Johann Rische, Martin Baumdick, Ana Jordan-Paiz, Julia Jäger, Ole Hinrichs, Deborah Sandfort (Department of Virus Immunology, Leibniz Institute of Virology; III. Department of Medicine and Hamburg Center for Translational Immunology, University Medical Center Hamburg-Eppendorf, Hamburg, Germany); Jakob Malsy (Department of Virus Immunology, Leibniz Institute of Virology; I. Department of Medicine, III. Department of Medicine and Hamburg Center for Translational Immunology, University Medical Center Hamburg-Eppendorf, Hamburg, Germany); Konrad Reinshagen, Christian Tomuschat (Department of Paediatric Surgery, University Medical Center Hamburg-Eppendorf, Hamburg, Germany); Nathaniel Melling (Department of General, Visceral and Thoracic Surgery, University Medical Center Hamburg-Eppendorf, Hamburg, Germany); and Ingo Königs (Department of Pediatric Surgery, University Medical Center Hamburg-Eppendorf; Department of Paediatric Surgery, Burn Unit, Plastic and Reconstructive Surgery, Altona Children’s Hospital, Hamburg, Germany). We also thank Jana Hennesen and Arne Düsedau from the flow cytometry core facility at the Leibniz Institute of Virology. We thank Sven Heins and Vadim Ustinov from the Institute of Medical Systems Biology (IMSB) at the University Medical Center Hamburg-Eppendorf for their IT support. We thank everyone from the IMSB at the University Medical Center Hamburg-Eppendorf and the Virus Immunology Research Group at Leibniz Institute of Virology who contributed with insightful discussions. This work was supported by the Daisy Hüet Roëll Foundation, the Landesforschungsförderung (LFF-75 and LFF-73) Hamburg City of Hamburg, the Deutsche Forschungsgemeinschaft (DFG) SFB1192, the SFB1328, BU3630/3-1 and EFRE 2014-2020 REACT-EU. This study is supported by the Innovative Antiviral Therapy Program, Leibniz Institute of Virology (LIV). The Leibniz Institute of Virology is supported by the Free and Hanseatic City of Hamburg and the Federal Ministry of Health, Germany. S.B. was supported by DFG SFB 1286 project Z2 and FOR 5068 project P9.

## Author contributions

M.B., S.B., and M.J.B. contributed to conceptualization and design. M.B. analyzed the data and developed the Seumetry R toolkit. M.F. collected and preprocessed the data. M.B., S.B., M.F., and M.J.B. drafted and revised the manuscript. Hamburg Intestinal Tissue Study Group was involved in the collection and processing of tissues and reviewed the manuscript.

## Supplementary tables

Table S1. Intestinal tissue samples.

Table S2. Surface proteins measured in intestinal flow cytometry data.

Table S3. Antibodies used for flow cytometry staining.

## Supplementary figures

**Figure S1.**
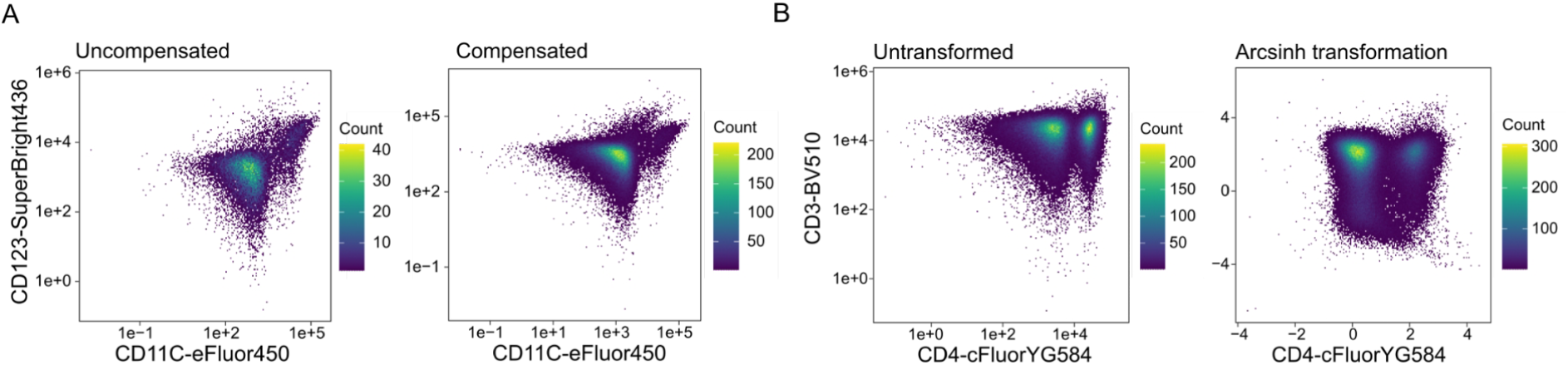
Compensation and transformation. (A) Representative two-dimensional density plots before and after compensation using CD11C and CD123 as exemplary markers (n=14). (B) Representative two-dimensional density plots of untransformed and inverse hyperbolic sine function (arcsinh) transformed data using CD3 and CD4 as exemplary markers (n=14).

**Figure S2.**
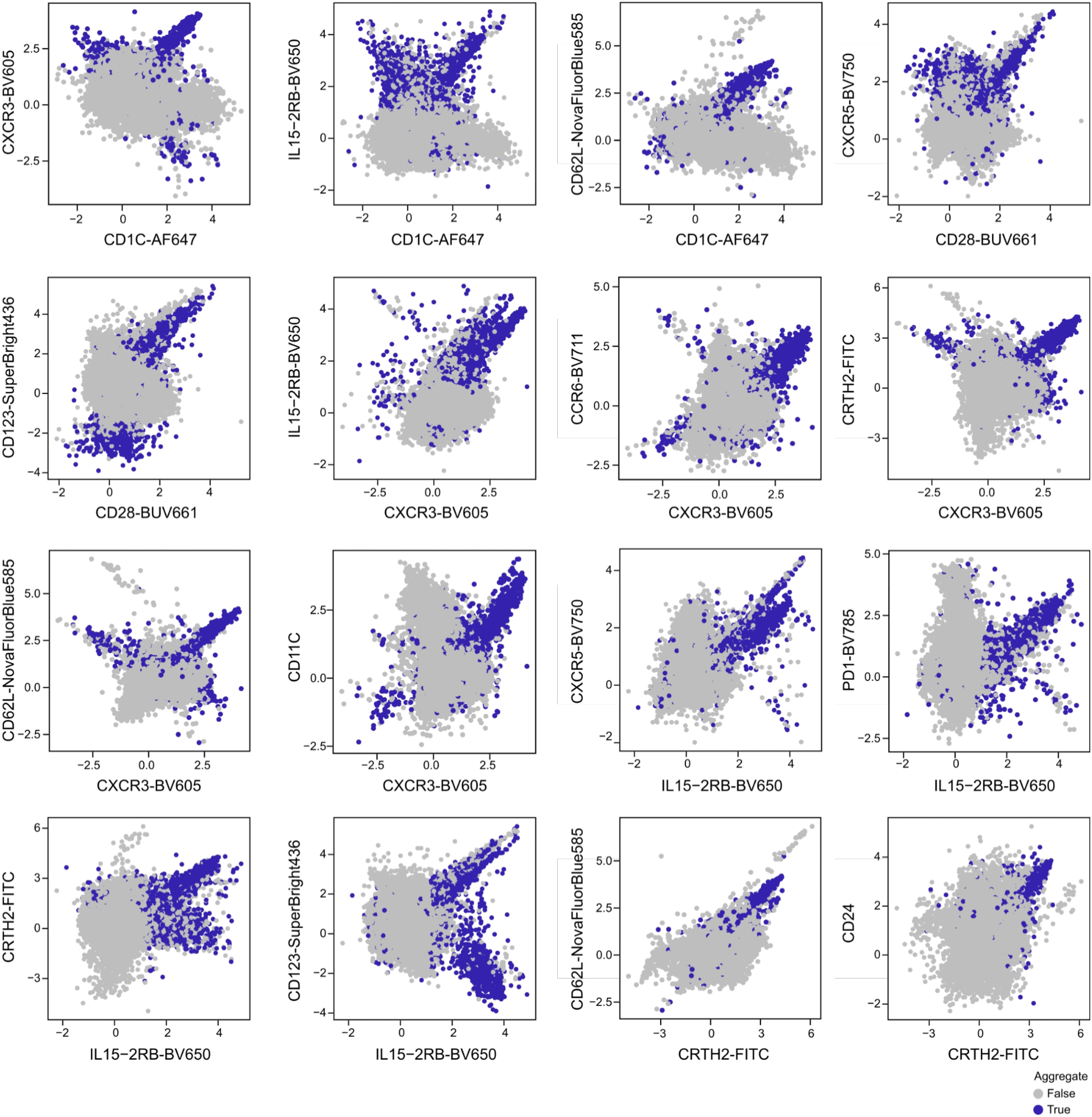
Antibody aggregate events across all channels. Antibody aggregates are detected using our novel algorithm on all channel combinations with a high likelihood of antibody aggregates, which are identified by Pearson correlation. Scatterplots depict identified antibody aggregates across all channels (blue) and non-aggregates (gray).

**Figure S3.**
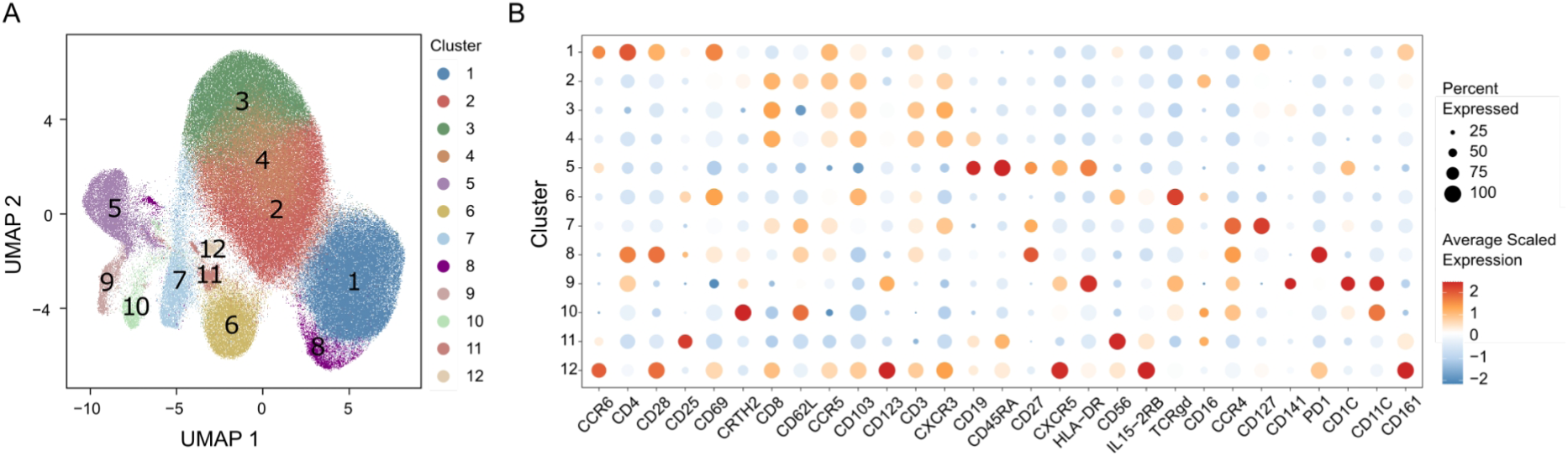
Merging and annotation of Louvain clusters identifying intestinal immune cell populations. (A) UMAP of all samples colored by Louvain clusters. (B) Dotplot of average expression (scaled) of top 5 cluster markers per cluster identified using Seurat’s “FindAllMarkers” function (n=14).

**Figure S4.**
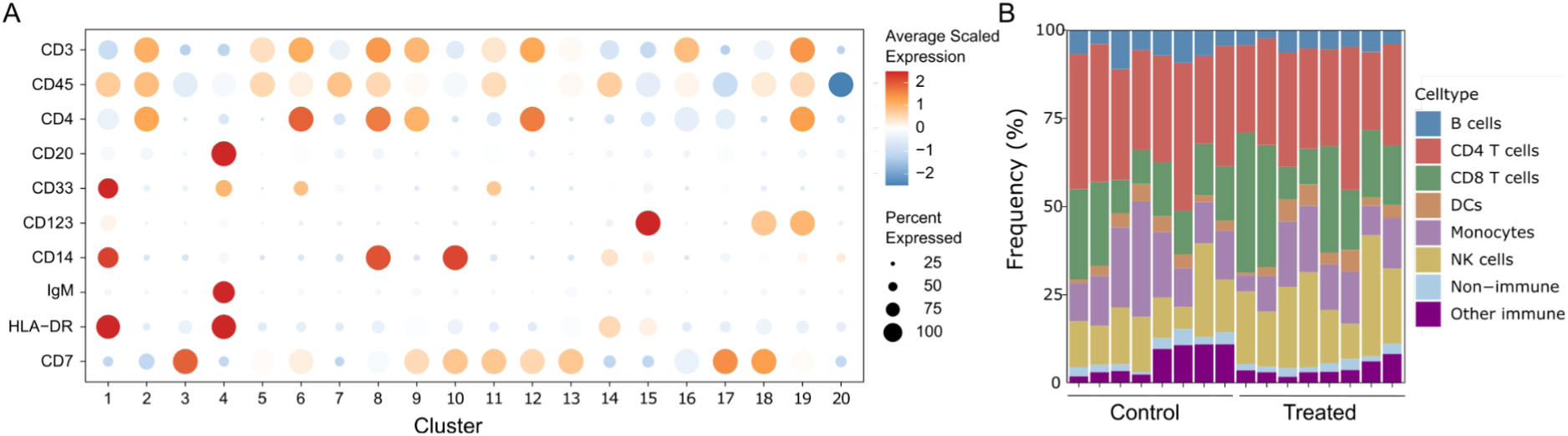
Expression of cluster markers and frequencies of immune cell populations in public dataset. (A) Dotplot of average lineage marker expression (scaled) in all 20 Louvain clusters. (B) Frequency plot of clusters across samples.

